# Neuronal KGB-1 JNK MAPK signaling regulates the dauer developmental decision in response to environmental stress in *C. elegans*

**DOI:** 10.1101/2021.06.28.450206

**Authors:** Deepshikha Dogra, Warakorn Kulalert, Frank C. Schroeder, Dennis H. Kim

## Abstract

In response to stressful growth conditions of high population density, food scarcity and elevated temperature, young larvae of nematode *Caenorhabditis elegans* can enter a developmentally arrested stage called dauer that is characterized by dramatic anatomic and metabolic remodeling. Genetic analysis of dauer formation of *C. elegans* has served as an experimental paradigm for the identification and characterization of conserved neuroendocrine signaling pathways. Here, we report the identification and characterization of a conserved JNK-like mitogen-activated protein kinase (MAPK) pathway that is required for dauer formation in response to environmental stressors. We observed that loss-of-function mutations in the MLK-1-MEK-1-KGB-1 MAPK pathway suppress dauer entry. Loss-of-function mutation in the VHP-1 MAPK phosphatase, a known negative regulator of KGB-1 signaling, results in constitutive dauer formation which is dependent on the presence of dauer pheromone but independent of diminished food levels or elevated temperatures. Our data suggest that KGB-1 pathway acts in the sensory neurons, in parallel to established insulin and TGF-*β* signaling pathways, to transduce the dauer-inducing environmental cues of diminished food levels and elevated temperature.

## Introduction

*Caenorhabditis elegans* can enter an alternative developmental diapause state, known as dauer, in response to unfavorable environmental conditions—specifically increased population density, scarcity of bacterial food, and elevated temperature (Golden and Riddle 1982; 1984a; 1984b; 1984c). The dauer state is characterized by increased stress resistance and dramatic remodeling of anatomy and metabolism (Cassada and Russell 1975). The genetic analysis of dauer formation has served as an experimental paradigm for understanding how neuroendocrine signaling through conserved components such as the insulin, TGF-*β*, and nuclear hormone receptor control organismal physiology and modulate developmental plasticity (Riddle and Albert 1997; Hu 2007; Fielenbach and Antebi 2008; Baugh and Hu 2020).

Recent studies have defined the molecular basis for how *C. elegans* responds to increased population density through pheromone signaling. Dauer pheromone has been characterized as a mixture of ascarosides, several structurally related derivatives of the dideoxysugar ascarylose (Butcher *et al.* 2007), which act on diverse families of G-protein-coupled membrane receptors. Receptors encoded by *srbc-64* and *srbc-66* are involved in perception of ascarosides ascr#1, ascr#2 and ascr#3, with probable role of additional GPCRs as the double mutant *srbc-64;srbc-66* still forms dauer at high concentrations of these ascarosides (Kim *et al.* 2009). McGrath *et al.* (2011) identified two predicted GPCRs, *srg-36* and *srg-37*, as ascr#5-specific receptors. Two additional receptors, DAF-37 and DAF-38, respond to ascr#2, ascr#3 and ascr#5 (Park *et al.* 2012). Pheromone signaling also modulates expression of DAF-7 (TGF-*β* ligand) and insulin ligands (Larsen, Albert and Riddle 1995; Ren *et al.* 1996; Schackwitz, Inoue and Thomas 1996; Kimura *et al.* 1997; Li, Kennedy and Ruvkun 2003), linking the activity of pheromone to canonical signaling pathways involved in the control of the dauer developmental decision.

In contrast, the mechanisms by which diminished levels of high-quality bacterial food or temperature can modulate the dauer decision remain less well understood. Food signal was first described as a compound present in bacterial extract that can inhibit dauer formation and initiate dauer recovery (Golden and Riddle 1982; 1984a). Bacterial fatty acids have been implicated in promoting dauer recovery (Kaul *et al.* 2014). Of note, recent studies have shown a role of Rictor/TORC2 and CaMKI in regulating expression of TGF-*β* ligand and insulin-like proteins in response to food signals in dauer entry (Neal *et al.* 2015; O’Donnell *et al.* 2018).

We previously described a genetic strategy to identify novel genes involved in dauer formation by undertaking a screen for suppressors of the *daf-28(sa191)* mutation (Kulalert and Kim 2013; Kulalert *et al.* 2017), which confers constitutive entry into dauer diapause (Malone, Inoue and Thomas 1996; Li, Kennedy and Ruvkun 2003). Here, we have identified and characterized a c-Jun N-terminal Kinase (JNK)-like mitogen-activated protein kinase (MAPK) KGB-1 pathway that functions in the sensory neurons to promote dauer formation. We characterize the effect of loss of the KGB-1 pathway, as well as its hyperactivation in a VHP-1 MAPK phosphatase mutant, on the regulation of the dauer developmental decision. Our data suggest a role for KGB-1 signaling in transducing diminished bacterial food and elevated temperature cues, which are integrated with pheromone signaling to promote entry into dauer diapause.

## Results

We previously reported the results of a genetic screen for suppressors of the dauer-constitutive (Daf-c) phenotype of the *daf-28(sa191)* mutant (Kulalert *et al.* 2017). Incomplete suppression of the Daf-c phenotype of the *daf-28(sa191)* mutant by *daf-3* and *daf-16* mutations (Malone, Inoue and Thomas 1996; Li, Kennedy and Ruvkun 2003; Kulalert and Kim 2013) suggested that genetic suppressors of *daf-28(sa191)* might uncover additional novel pathways functioning in parallel to established signaling pathways activating the dauer developmental decision. Our prior characterization of the molecular mechanisms driving dauer entry of the *daf-28(sa191)* mutant determined that activation of the Unfolded Protein Response and PEK-1 phosphorylation of eIF2*α* in the ASI neuron pair conferred the Daf-c phenotype (Kulalert and Kim 2013; Kulalert *et al.* 2017). We reported the isolation of multiple suppressor mutations in genes acting in this pathway downstream of UPR activation in the ASI neuron pair (Kulalert *et al.* 2017). We also reasoned that we might isolate suppressor mutations in novel pathways acting in parallel to this and previously characterized pathways leading to dauer formation. We recovered a strain carrying the *qd318* mutation from a genetic screen for suppressors of the Daf-c phenotype of the *daf-28(sa191)* mutant (Kulalert *et al.* 2017). Molecular characterization of this strain revealed a mutation in the *mlk-1* gene encoding a mitogen-activated protein (MAP) kinase kinase kinase. We sought to determine whether MLK-1 is also required for dauer entry in wild-type animals in the absence of the *daf-28(sa191)* trigger for dauer entry.

Induction of dauer formation under experimental conditions involves the addition of dauer pheromone, which is a combination of ascaroside molecules produced by *C. elegans* (Jeong *et al.* 2005; Butcher *et al.* 2007; Butcher *et al.* 2008; Pungaliya *et al.* 2009; Gallo and Riddle 2009), reduction of high-quality bacterial food *E. coli* OP50, and incubation at higher temperatures (Golden and Riddle 1984a; 1984b; Ailion and Thomas 2000; Neal, Kim and Sengupta 2013). Figure 1a depicts an overview of the environmental conditions that affect the decision to enter dauer. We experimentally imposed environmental stressors of crowding, low food availability and elevated ambient temperature in the laboratory, by using a combination of pheromone, heat-killed *E. coli* OP50 and 25°C (Figure 1b). Under these conditions, we observed that > 80% of wild-type larvae formed dauers (Figure 1c). In contrast, we observed that dauer formation was largely blocked in the *mlk-1(qd318)* mutant and a mutant carrying a deletion in the *mlk-1* gene, *mlk-1(km19)*.

**Figure 1.**
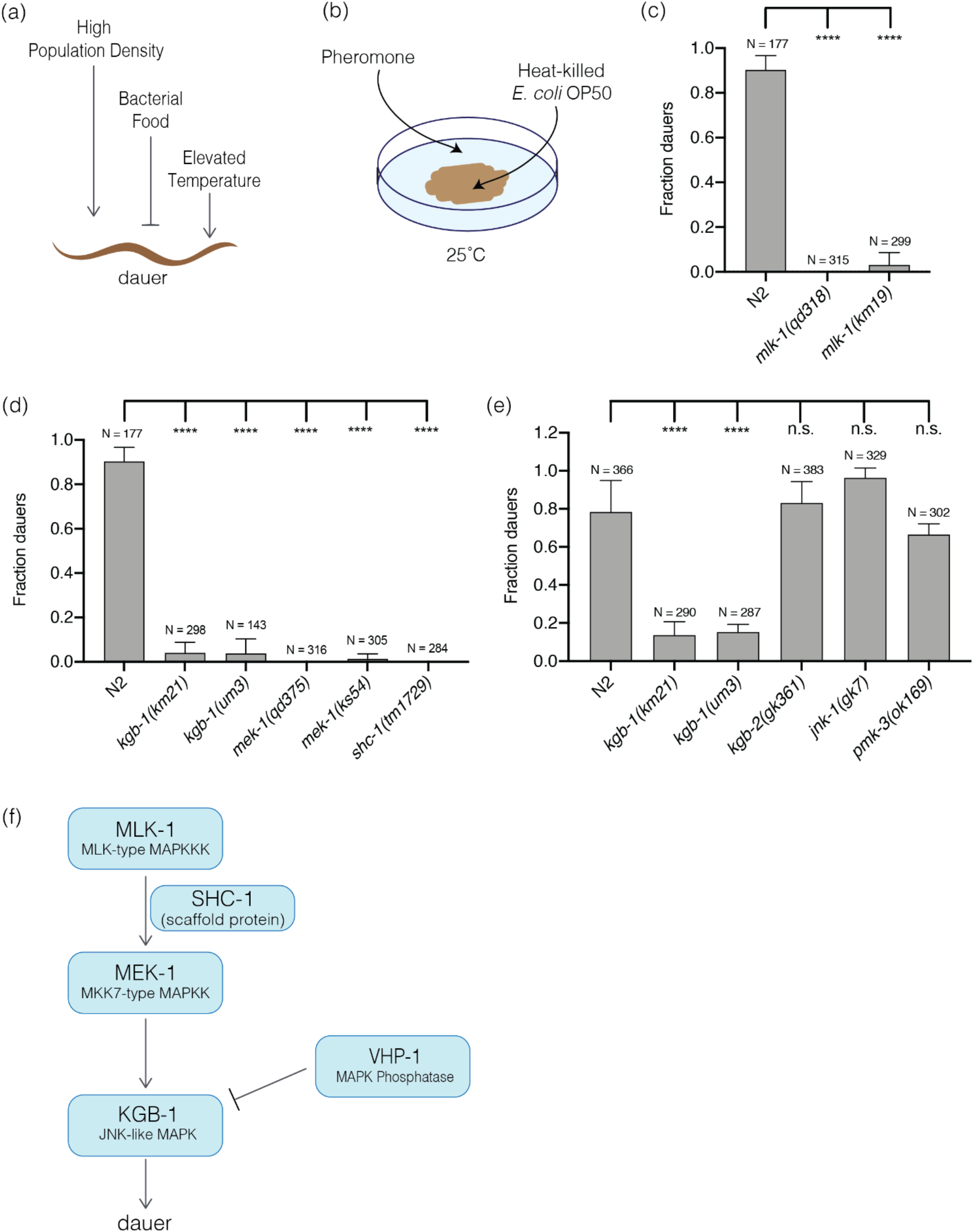
KGB-1 pathway activation is required for dauer entry caused by environmental stress. (a) Schematic depiction of environmental conditions that promote and suppress dauer formation. High population density and elevated temperature promote (shown by pointing arrows), and abundant food supply suppresses (shown by inhibitory arrow) dauer formation. (b) Schematic of dauer inducing conditions: a combination of pheromone, heat-killed *E. coli* OP50 and 25°C, applied to noble agar plates in lab to emulate the environmental conditions promoting dauer formation depicted in (a). (c) Fraction of wild-type (N2) animals, and *mlk-1(qd318)* and *mlk-1(km19)* mutant animals entering dauer in the presence of dauer inducing conditions. Statistical analysis conducted with ordinary one-way ANOVA followed by Dunnett’s multiple comparisons test. Plotted is mean + SD, N = total number of animals tested. (d) Fraction of wild-type (N2) animals, and *kgb-1(km21), kgb-1(um3), mek-1(qd375), mek-1(ks54)* and *shc-1(tm1729)* mutant animals entering dauer in the presence of dauer-inducing conditions. Statistical analysis conducted with ordinary one-way ANOVA followed by Dunnett’s multiple comparisons test. Plotted is mean + SD, N = total number of animals tested. (e) Fraction of wild-type (N2) animals, and *kgb-1(km21), kgb-1(um3), kgb-2(gk361), jnk-1(gk7)* and *pmk-3(ok169)* mutant animals entering dauer in the presence of dauer inducing conditions. Statistical analysis conducted with ordinary one-way ANOVA followed by Dunnett’s multiple comparisons test. Plotted is mean + SD, N = total number of animals tested, n.s. = not significant. (f) Schematic of KGB-1 pathway, a c-Jun N-terminal kinase (JNK)-like mitogen activated protein kinase pathway, composed of MLK-1 (MLK-type MAP3K), MEK-1 (MKK7-type MAP2K), SHC-1 (scaffold protein), KGB-1 (JNK-like MAPK) and VHP-1 (MAPK phosphatase).

MLK-1 has been previously characterized as acting upstream of the KGB-1 MAPK, which is homologous to the conserved c-Jun-N-terminal Kinase (JNK) MAPK. MLK-1 phosphorylates the MAPK kinase MEK-1, a homolog of the MKK7 MAPKK, in the presence of the scaffold protein SHC-1, and MEK-1 phosphorylates and activates KGB-1 (Sakaguchi, Matsumoto and Hisamoto 2004; Mizuno *et al.* 2004; Mizuno *et al.* 2008). All the kinase proteins in this pathway as well as the scaffold protein are essential for activation of KGB-1. The animals with loss-of-function mutations in genes encoding components of the KGB-1 pathway continued their reproductive growth, failing to enter dauer under these stressful conditions (Figure 1d). We determined that the KGB-1 MAPK is the only JNK MAPK required for dauer entry, as only *kgb-1* loss-of-function mutants (*km21* and *um3*) suppressed dauer entry, whereas the animals carrying mutations in the other two JNK homologs, *jnk-1(gk7)* and *kgb-2(gk361),* were able to enter dauer similar to wild-type animals in the presence of the dauer inducing conditions (Figure 1e). The KGB-1 pathway was reported to have cross-talk with PMK-3 (p38 MAPK) pathway in axon regeneration where both the pathways are required (Nix *et al.* 2011), but we observed that the *pmk-3* loss-of-function mutant exhibited wild-type dauer entry (Figure 1e). Our data establish that the MLK-1-(SHC-1)-MEK-1-KGB-1 JNK MAPK pathway is required for dauer entry (Figure 1f).

To determine the tissue and cell types in which the KGB-1 pathway functions to promote dauer entry, we conducted tissue-specific rescue experiments of the *mek-1(qd375)* mutant using transgenes expressing the *mek-1* cDNA under the control of heterologous promoters. We fully rescued the Daf-d phenotype of the *mek-1(qd375)* mutant when expressing *mek-1* under its endogenous promoter and 3’UTR in an extrachromosomal array (Figure 2a). We observed that expression of *mek-1* in the nervous system of *mek-1(qd375)* mutant animals, under the control of the pan-neuronal promoter, *rab-3p*, resulted in rescue of the dauer formation phenotype, whereas expression of *mek-1* in the intestine or body wall muscles failed to rescue the Daf-d phenotype of the *mek-1(qd375)* mutant (Figure 2b-d). In particular, we introduced expression of *mek-1* under the *bbs-1p* promoter, which directs expression in all ciliated neurons, and observed that expression in this specific subset of neurons rescued the Daf-d phenotype of *mek-1(qd375)* mutant animals (Figure 2e). Ciliated neurons encompass the chemosensory nervous system of *C. elegans*, and a number of specific sensory neurons have been implicated in roles in entry into, and exit from, dauer diapause (Albert, Brown and Riddle 1981; Bargmann and Horvitz 1991; Schackwitz, Inoue and Thomas 1996; Ren *et al.* 1996; Li, Kennedy and Ruvkun 2003; Bargmann 2006; Ludewig and Schroeder 2018), but we were unable to further define specific neurons involved in dauer entry using neuronal cell type-specific rescue experiments. These data suggest either limitations of our experimental approach or the function of KGB-1 across multiple ciliated chemosensory neurons in dauer formation.

**Figure 2.**
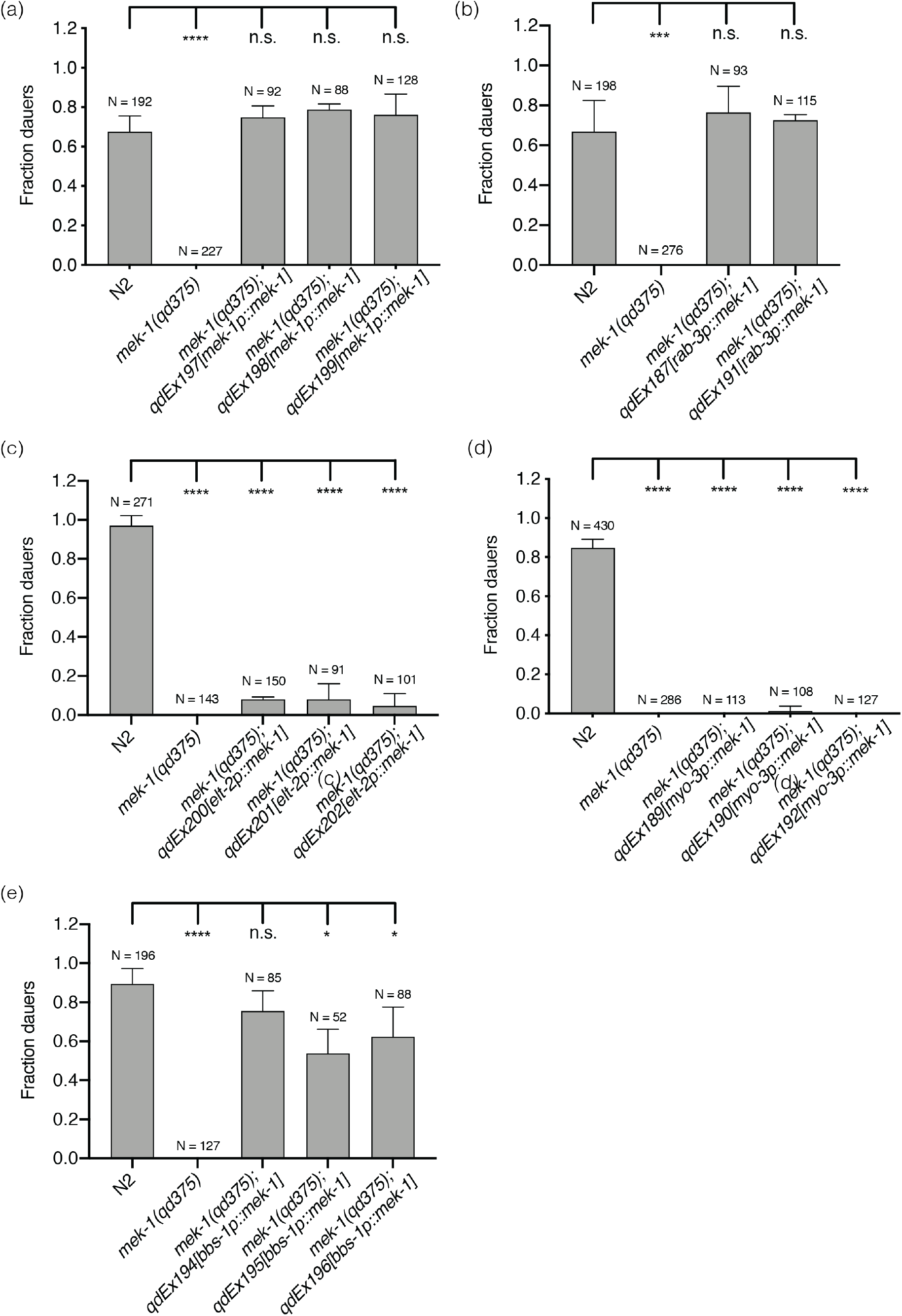
KGB-1 pathway acts in the chemosensory neurons to regulate dauer diapause. Fraction of wild-type (N2) animals, *mek-1(qd375)* mutant animals (non-sibling control) and *mek-1(qd375)* animals with extrachromosomal expression of (a) endogenous *mek-1* gene (b) *mek-1* gene under pan-neuronal promoter, *rab-3p* (c) *mek-1* gene under intestine-specific promoter, *elt-2p* (d) *mek-1* gene under body-wall muscle specific promoter, *myo-3p* (e) *mek-1* gene under pan-ciliated neuron specific promoter, *bbs-1p*, entering dauer in the presence of pheromone and heat-killed *E. coli* OP50 at 25°C. Statistical analysis conducted with ordinary one-way ANOVA followed by Dunnett’s multiple comparisons test. Plotted is mean + SD, N = total number of animals tested, n.s. = not significant.

Having observed the effect of loss of KGB-1 signaling on blocking entry into dauer diapause, we sought to examine if increased KGB-1 signaling would confer the opposite phenotype of enhancing dauer formation. The VHP-1 MAPK phosphatase was previously characterized as a negative regulator of the KGB-1 MAPK, and diminished activity of VHP-1 resulted in hyperactivation of KGB-1 (Mizuno *et al.* 2004). A strain carrying a *vhp-1(km20)* deletion was previously noted to exhibit early larval lethality (Mizuno *et al.* 2004), but a viable *vhp-1* mutant carrying a nonsense mutation, *vhp-1(sa366)* has also been reported (Choy and Thomas 1999), and thus we examined this mutant for a dauer formation phenotype.

We examined wild-type and mutant larvae under twelve different sets of conditions derived from combinations of the presence or absence of pheromone, the presence of high quality live bacterial food or poor quality heat-killed bacterial food, and three temperatures (25°C, 20°C, 16°C). The experimental data are presented in a 2×2 array, with each cell distinguished by the presence or absence of pheromone and whether larvae were exposed to live bacterial food or heat-killed bacterial food. Within each cell, the results of the three different incubation temperatures are presented. As we indicated above, optimal induction of dauer formation in wild-type animals was observed in the presence of dauer pheromone and heat-killed bacterial food at the elevated temperature of 25°C (Figure 3a). We did not observe dauer formation in the *vhp-1(sa366)* mutant in the absence of pheromone, even under conditions of bacterial food deprivation and elevated temperature, just as we observed for wild-type larvae. In the presence of pheromone and heat-killed food, we observed high-level dauer formation in *vhp-1(sa366)* larvae as was observed for wild-type animals. However, in contrast to what we observed for wild-type larvae, we detected equivalent high levels of dauer entry even at the lower temperatures of 20°C and 16°C under these conditions (Figure 3b). More dramatic differences were observed under conditions in the presence of pheromone and live bacterial food. In wild-type animals, the presence of live bacterial food largely suppressed dauer formation, but in *vhp-1(sa366)* animals, we observed marked dauer entry, equivalent to levels observed in the presence of heat-killed bacterial food.

**Figure 3.**
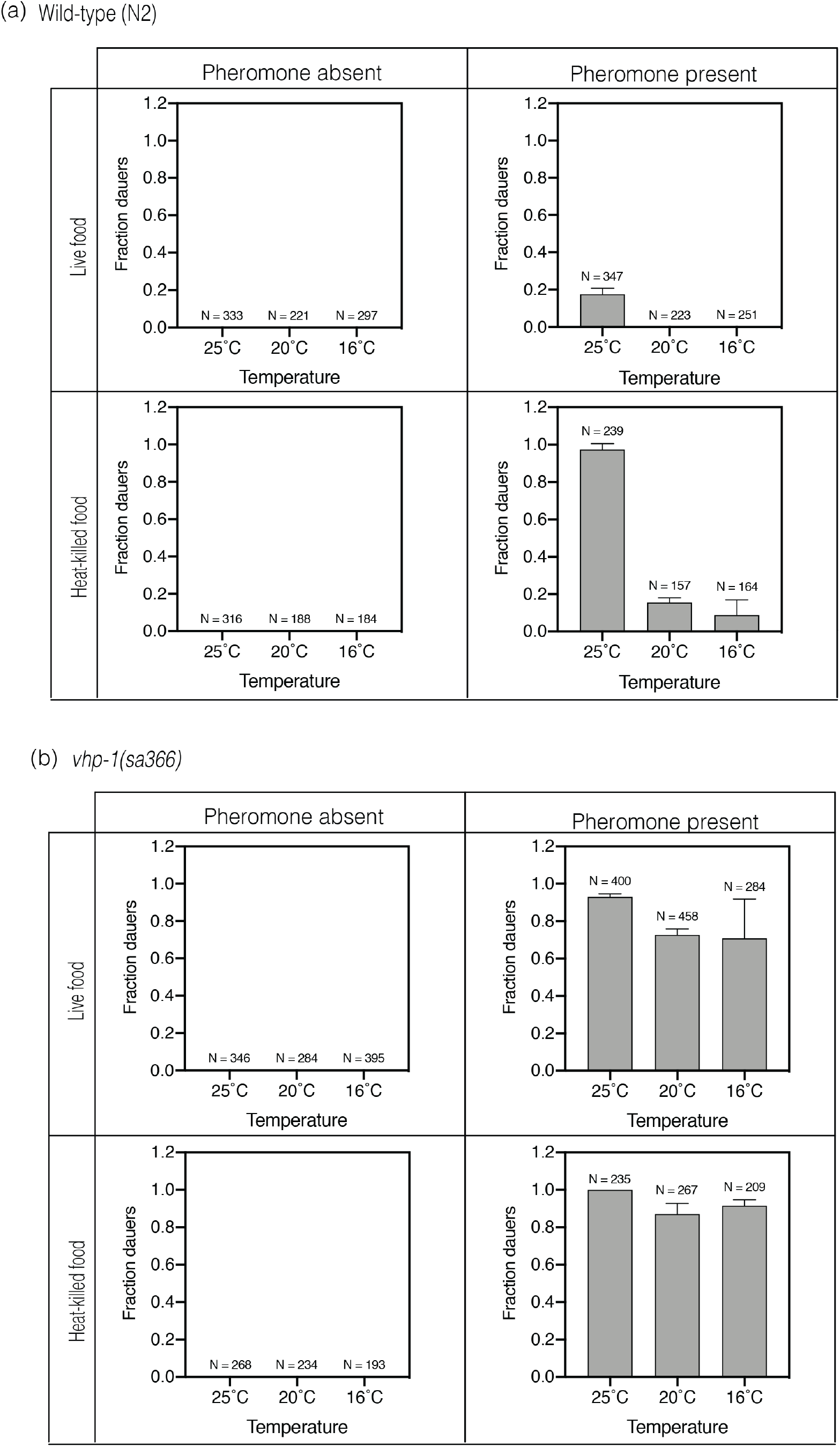
Increased activation of the KGB-1 pathway in the *vhp-1* mutant leads to constitutive dauer formation in the presence of pheromone. Fraction of (a) wild-type animals (b) *vhp-1(sa366)* mutant animals entering dauer under various combinations of dauer inducing conditions. The first row shows fraction dauers in the presence of live *E. coli* OP50 and the second row shows fraction dauers in the presence of heat-killed *E. coli* OP50. The first column shows fraction dauers in the absence of pheromone and the second column shows fraction dauers in the presence of pheromone. Each graph in the four panels shows fraction dauers at 25C, 20°C and 16°C respectively. Plotted is mean + SD, N = total number of animals tested.

We confirmed that *vhp-1(sa366)* is a hypomorphic allele by genetic analysis of trans-heterozygote animals using wild-type animals and the *vhp-1(km20)* null mutant (Figure 4a-d). We also confirmed that dauer formation in the *vhp-1(sa366)* mutant was suppressed by mutation in the KGB-1 pathway, specifically a mutation in the *mek-1* gene encoding the MAPKK MEK-1 that phosphorylates KGB-1 (Figure 4e-f). In the absence of phosphorylation of KGB-1, reduction of function of the VHP-1 phosphatase would not be expected to affect KGB-1 activity.

**Figure 4.**
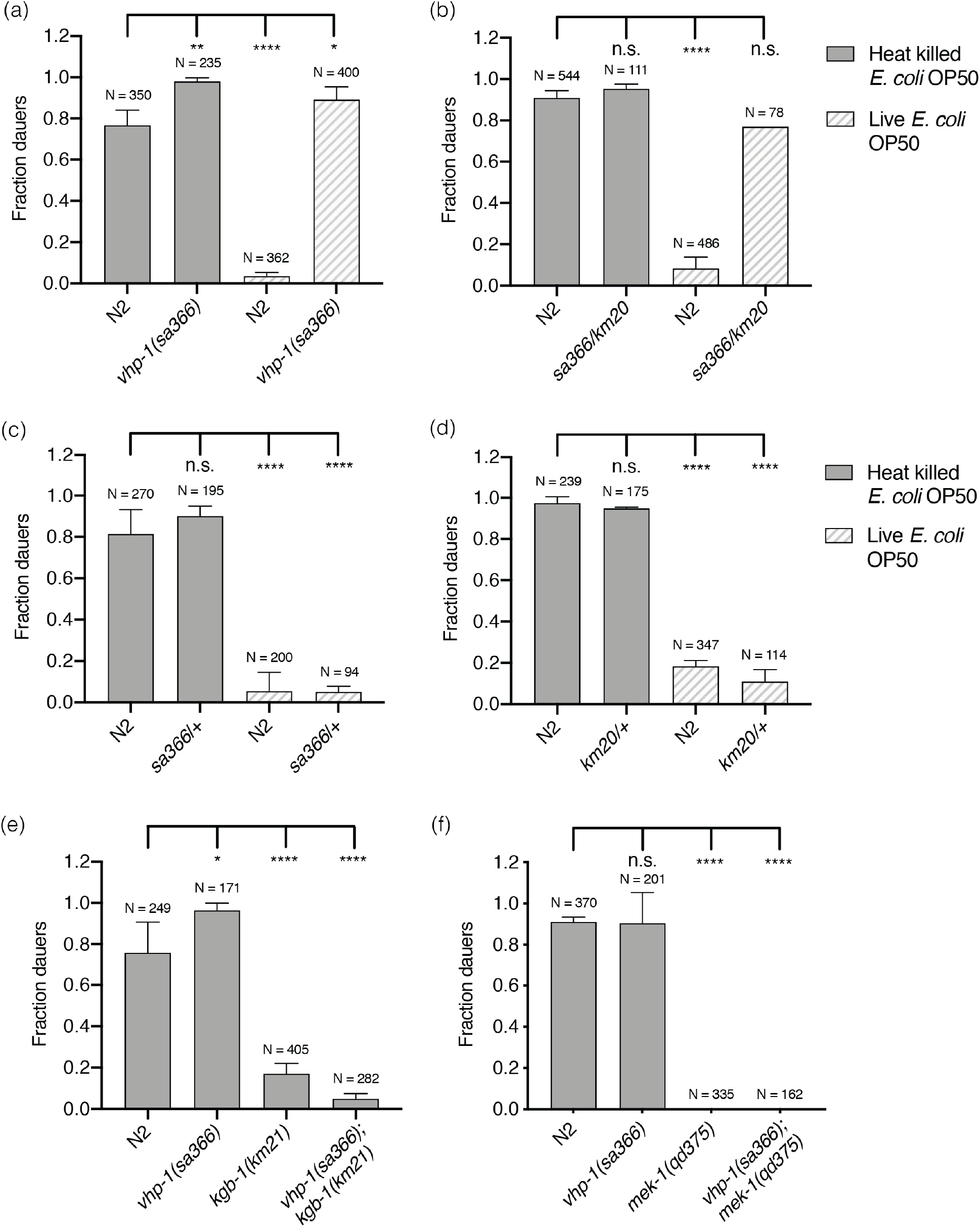
Dauer formation of *vhp-1(sa366)* mutant animals is due to hyperactivation of KGB-1 pathway. (a-d) Fraction of wild-type (N2) animals and (a) *vhp-1(sa366)* homozygous mutant animals (b) *vhp-1(sa366/km20)* heterozygous mutant animals (c) *vhp-1(sa366/+)* heterozygous mutant animals (d) *vhp-1(km20/+)* heterozygous mutant animals, entering dauer in the presence of pheromone, 25°C and either heat-killed (solid bars) or live (patterned bars) *E. coli* OP50. Statistical analysis conducted with ordinary one-way ANOVA followed by Dunnett’s multiple comparisons test. Plotted is mean + SD, N = total number of animals tested, n.s. = not significant. (e-f) Fraction of wild-type (N2) animals, *vhp-1(sa366),* (e) *kgb-1(km21)* and *vhp-1(sa366);kgb-1(km21),* (f) *mek-1(qd375)* and *vhp-1(sa366);mek-1(qd375)* mutant animals, entering dauer in the presence of pheromone, 25°C and heat-killed *E. coli* OP50. Statistical analysis conducted with ordinary one-way ANOVA followed by Dunnett’s multiple comparisons test. Plotted is mean + SD, N = total number of animals tested, n.s. = not significant.

These data suggest that increased activity of the KGB-1 pathway in the *vhp-1(sa366)* mutant can substitute for the requirement for diminished bacterial food availability and elevated temperature in the induction of dauer formation, whereas the requirement for pheromone is unaffected by increased KGB-1 activity. We next sought to define the genetic interactions among KGB-1 signaling and established pathways involved in the dauer developmental decision. We determined that the enhanced dauer entry phenotype of *vhp-1(sa366)* mutant larvae was suppressed by *daf-12* (Figure 5a, 5b), as is the case with most known dauer-constitutive mutants (Vowels and Thomas 1992). We observed that the *vhp-1(sa366)*; *daf-16(mu86)* mutant in the presence of pheromone resulted in the formation of partial dauers, regardless of whether live or heat-killed bacterial food was present (Figure 5c, 5d). Partial dauer formation was also observed in double mutants involving *daf-16* and dauer-constitutive mutants, including *daf-28(sa191)* (Vowels and Thomas 1992; Malone, Inoue and Thomas 1996), but not observed in *daf-2;daf-16* double mutants. The observation of partial dauers in the *vhp-1(sa366);daf-16(mu86)* mutant animals suggests that the KGB-1 signaling can activate the developmental decision to commit to dauer entry and partially proceed independent of DAF-16, but that the full execution of dauer program requires DAF-16 activity.

**Figure 5.**
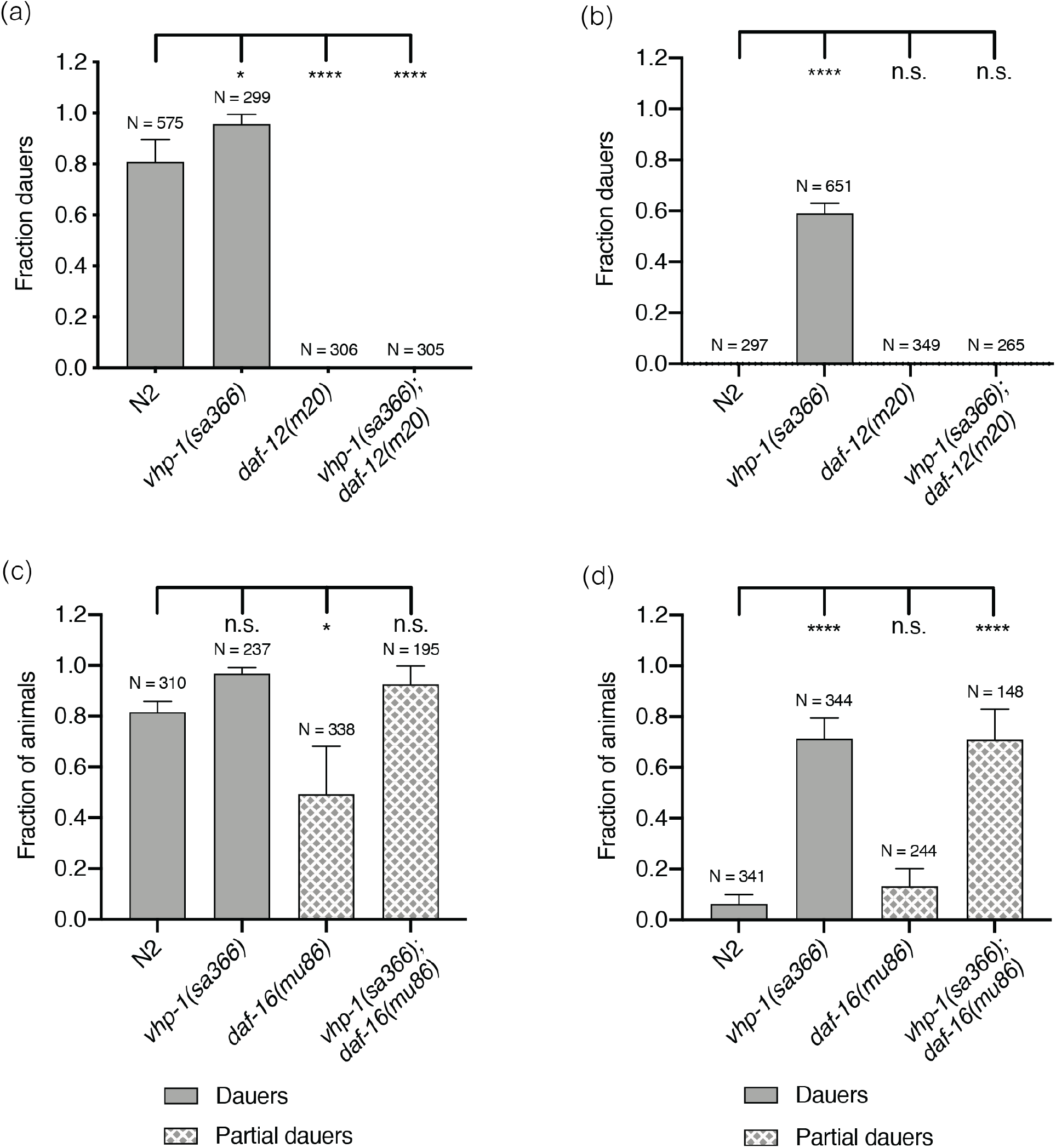
KGB-1 pathway functions upstream of the steroidal hormone pathway and in parallel to the insulin signaling pathway. (a-b) Fraction of wild-type (N2) animals and *vhp-1(sa366), daf-12(m20)* and *vhp-1(sa366);daf-12(m20)* mutant animals entering dauer in the presence of pheromone, 25°C and (a) heat-killed *E. coli* OP50 (b) live *E. coli* OP50. Statistical analysis conducted with ordinary one-way ANOVA followed by Dunnett’s multiple comparisons test. Plotted is mean + SD, N = total number of animals tested, n.s. = not significant. (c-d) Fraction of wild-type (N2) animals and *vhp-1(sa366), daf-16(mu86)* and *vhp-1(sa366);daf-16(mu86)* mutant animals entering dauer in the presence of pheromone, 25°C and (c) heat-killed *E. coli* OP50 (d) live *E. coli* OP50. Statistical analysis conducted with ordinary one-way ANOVA followed by Dunnett’s multiple comparisons test. Plotted is mean + SD, N = total number of animals tested, n.s. = not significant.

We observed that *vhp-1(sa366);daf-3(e1376)* mutants did not enter dauer in the presence of pheromone (Figure 6a, 6b). One interpretation of this observation was that KGB-1 signaling might function upstream of DAF-7-DAF-3 signaling. Alternatively, because pheromone is required for *vhp-1(sa366)*-enhanced dauer formation, we considered the possibility that KGB-1 signaling might function in parallel with DAF-7-DAF-3 signaling, with the *daf-3* mutation attenuating the effect of pheromone. Indeed, *daf-7* mutants exhibit a dauer-constitutive phenotype in the absence of pheromone, and pheromone downregulates *daf-7* expression in the ASI sensory neurons (Thomas, Birnby and Vowels 1993; Ren *et al.* 1996). To distinguish between these possibilities, we utilized a previously described reporter of DAF-3 activity, *cuIs5* (Thatcher *et al.* 1999; Reiner *et al.* 2008). This reporter consists of *myo-2* promoter, preceded by tandem copies of C183 portion of C-subelement, driving GFP expression in the pharynx. DAF-3 specifically recognizes and binds to C183 causing a suppression of GFP expression when it is not inhibited by DAF-7 (Figure 6c). As expected and has been reported, in the *daf-7* mutant background, DAF-3 reporter activity is diminished. The presence of pheromone also decreased DAF-3 reporter activity relative to conditions in the absence of pheromone although no difference was observed in the *daf-7* mutant, consistent with the *daf-7* mutant mimicking conditions of pheromone presence. If *vhp-1(sa366)* promoted dauer entry by modulating DAF-3 activity, we would expect DAF-3 reporter activity to be similarly diminished in the *vhp-1(sa366)* background in the presence of pheromone. We observed that DAF-3 reporter activity was not affected by the *vhp-1(sa366)* mutation (Figure 6d). These data suggest that the *vhp-1(sa366)* mutation and corresponding KGB-1 activity function in parallel to TGF-*β* signaling to promote dauer entry, and indicate that pheromone may act in part through modulation of the DAF-3-dependent signaling pathway.

**Figure 6.**
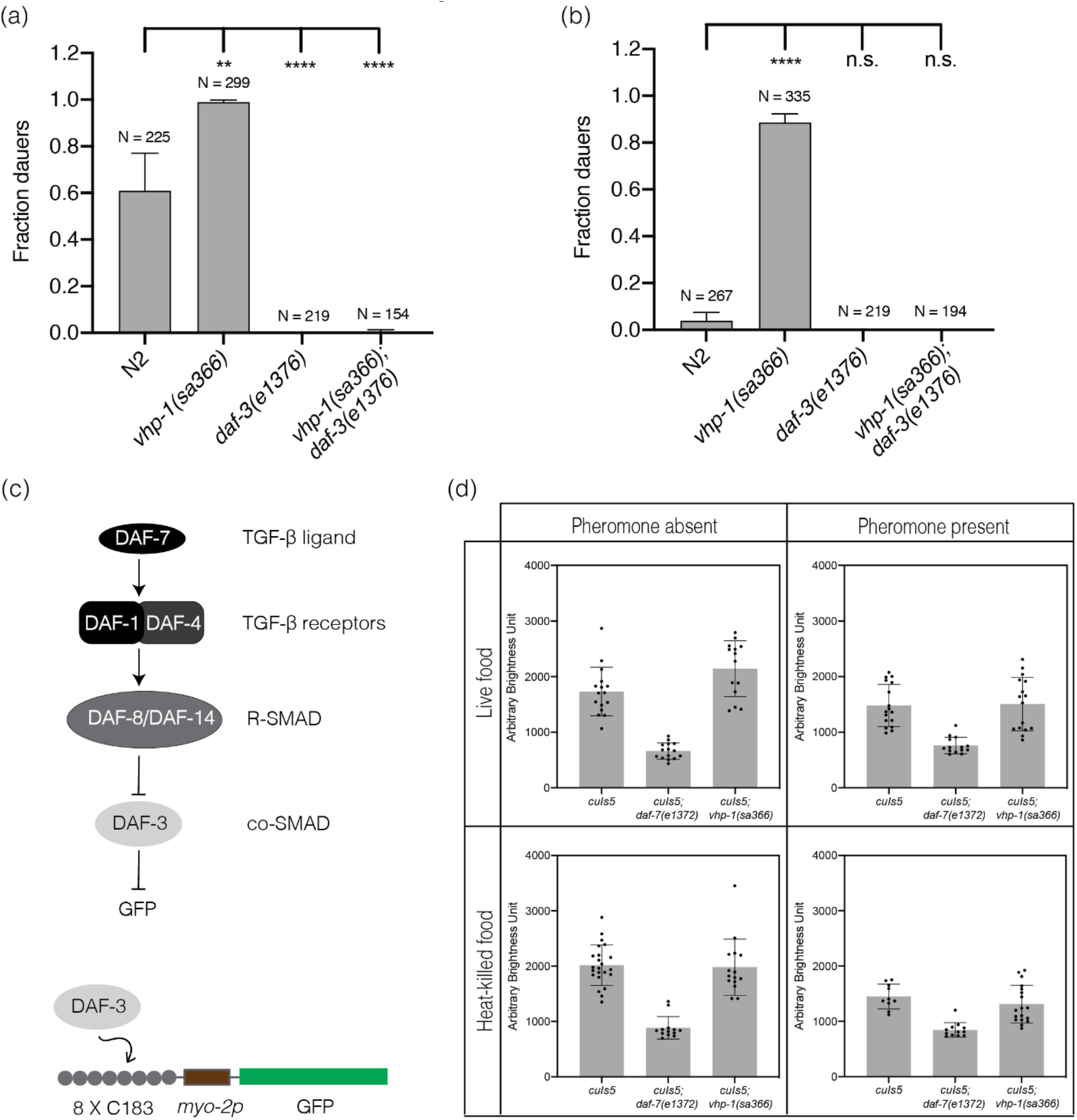
KGB-1 pathway functions in parallel to TGF-β signaling pathway in regulation of dauer diapause (a-b) Fraction of wild-type (N2) animals, *vhp-1(sa366), daf-3(e1376)* and *vhp-1(sa366);daf-3(e1376)* mutant animals entering dauer in the presence of pheromone, 25°C and (a) heat-killed *E. coli* OP50 (b) live *E. coli* OP50. Statistical analysis conducted with ordinary one-way ANOVA followed by Dunnett’s multiple comparisons test. Plotted is mean + SD, N = total number of animals tested, n.s. = not significant. (c) Overview of TGF-β signaling pathway and schematic of *cuIs5 -* DAF-3 reporter transgene. (d) Quantification of GFP expression from DAF-3 reporter (*cuIs5)*. Each graph in the four panels shows GFP expression in wild-type background, *daf-7(e1372)* and *vhp-1(sa366)* mutant backgrounds respectively. The first row shows fraction dauers in the presence of live *E. coli* OP50 and the second row shows fraction dauers in the presence of heat-killed *E. coli* OP50. The first column shows fraction dauers in the absence of pheromone and the second column shows fraction dauers in the presence of pheromone. Plotted is mean + SD, each dot represents an animal.

**Figure 7.**
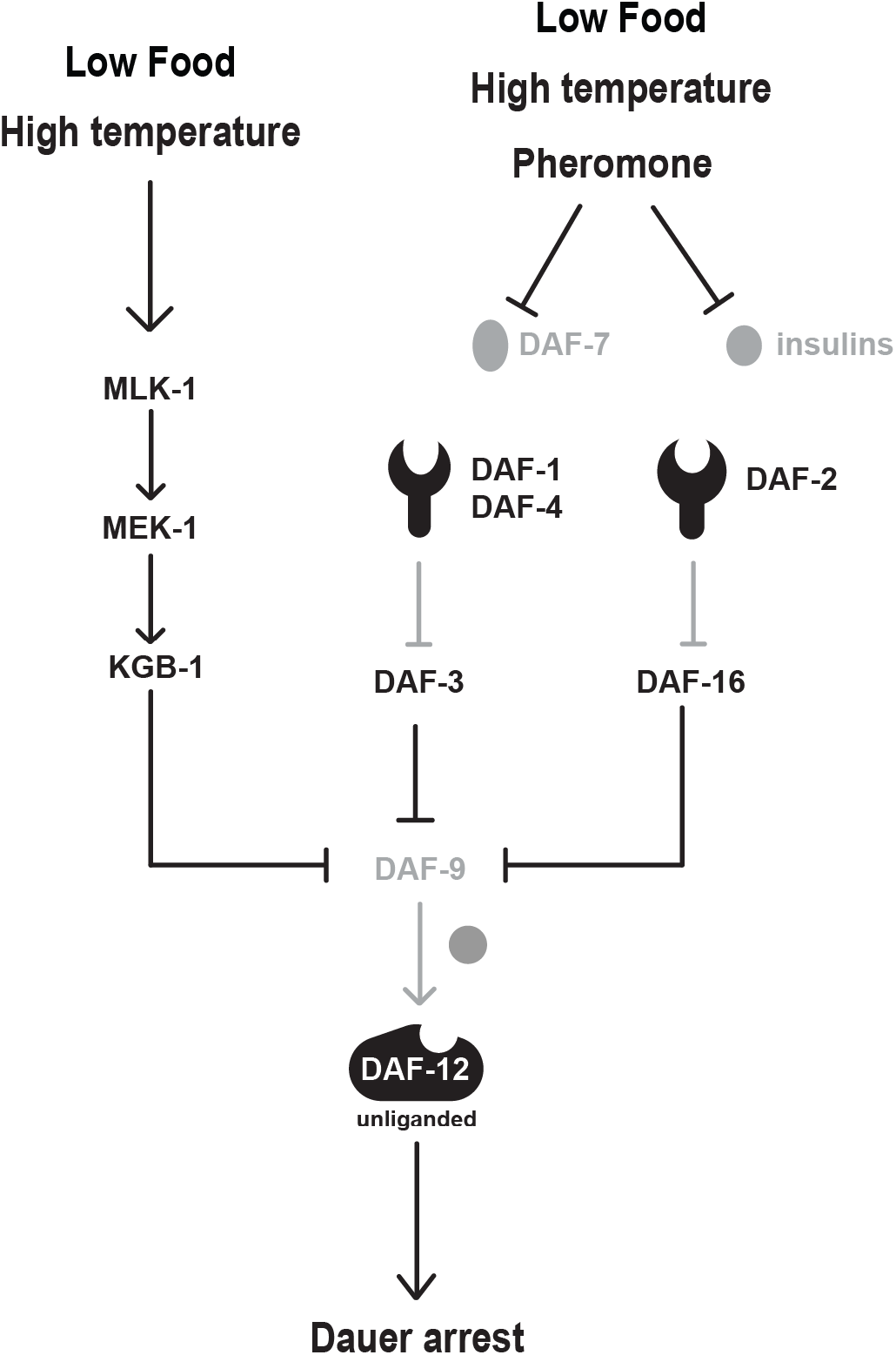
Model figure depicting KGB-1 pathway functioning parallel to TGF-β and insulin signaling pathways and upstream of steroid hormone pathway, in response to stress conditions of diminished food and elevated temperature to regulate dauer formation.

## Discussion

Dauer entry represents an integrated organismal response to environmental stress during development. Stress-activated MAPK signaling pathway have evolutionarily conserved roles in a wide range of stress responses in diverse species, and the KGB-1 JNK MAPK pathway has been implicated in the *C. elegans* response to multiple stressors, including heavy metals (Koga *et al.* 2000; Mizuno *et al.* 2004), the ER toxin tunicamycin (Mizuno *et al.* 2008), and food deprivation in the context of lifespan extension through intermittent fasting (Uno *et al.* 2013). KGB-1 is also required for resistance to pathogenic bacteria (Kim *et al.* 2004) and pore-forming toxins (Huffman *et al.* 2004). Here, our data suggest a role for KGB-1 signaling in the dauer developmental decision, with increased KGB-1 signaling in the *vhp-1* mutant enabling dauer formation independent of bacterial food availability and temperature. Previous studies establishing an age-dependent role of KGB-1 pathway described interaction with insulin signaling pathway, specifically DAF-16. (Twumasi-Boateng and Liu Shapira refs). Our epistasis analysis suggests that the KGB-1 pathway functions in parallel to insulin signaling in the decision to enter dauer diapause, but that full execution of the dauer program requires DAF-16 as indicated by the formation of incomplete, or partial dauers in the absence of DAF-16.

Early studies established that the ratio of pheromone to bacterial food is more important than the absolute concentration *per se* in regulating dauer entry. Using serial dilutions of both pheromone and food concentrations, Riddle and colleagues showed that dauer phenotype is dependent on the relative concentration of the signals (Golden and Riddle 1982; 1984a). Higher pheromone concentration induces a greater fraction of the population to enter dauer but this effect saturates at a fraction dependent on the total amount of bacterial food present (Golden and Riddle 1984b). Similarly, increases in temperature enhance dauer entry in the presence of a constant pheromone to food ratio (Golden and Riddle 1984b; 1984c). Our results suggest a role of the KGB-1 pathway in the interpretation of the pheromone to food ratio in the dauer developmental decision. Whereas the prior identification of dauer-defective mutants and, correspondingly, signaling pathways that promote dauer formation have largely been in the experimental context of suppression of Daf-c mutations, recent studies have focused on the genetic characterization of dauer entry in wild-type larvae in response to environmental cues. Of note, mutants of Rictor/TORC2 exhibit high temperature (27°C) induced dauer entry, in a manner that was also not affected by bacterial food quality, and the Rictor/TORC2 pathway was implicated in relaying food-sensing information from the intestine to the nervous system (O’Donnell *et al.* 2018). Whereas *rict-1* mutants modulated neuronal expression of *daf-7*, our data suggest that KGB-1 signaling mediates food sensitivity in parallel to the DAF-7*/*TGF-*β* signaling pathway pathway. Our study suggests that the activity of a conserved stress-activated KGB-1 signaling pathway in the sensory nervous system is required for the dauer developmental decision in response to pheromone and diminished food availability, functioning in parallel to established neuroendocrine pathways involved in the regulation of dauer formation.

## Materials and Methods

### Strains

All *C. elegans* strains used in this study were maintained as previously described (Brenner 1974). The temperature sensitive strains were maintained at 16 °C and all other strains were incubated at 20°C. The following is a complete list of all the strains used in this study:

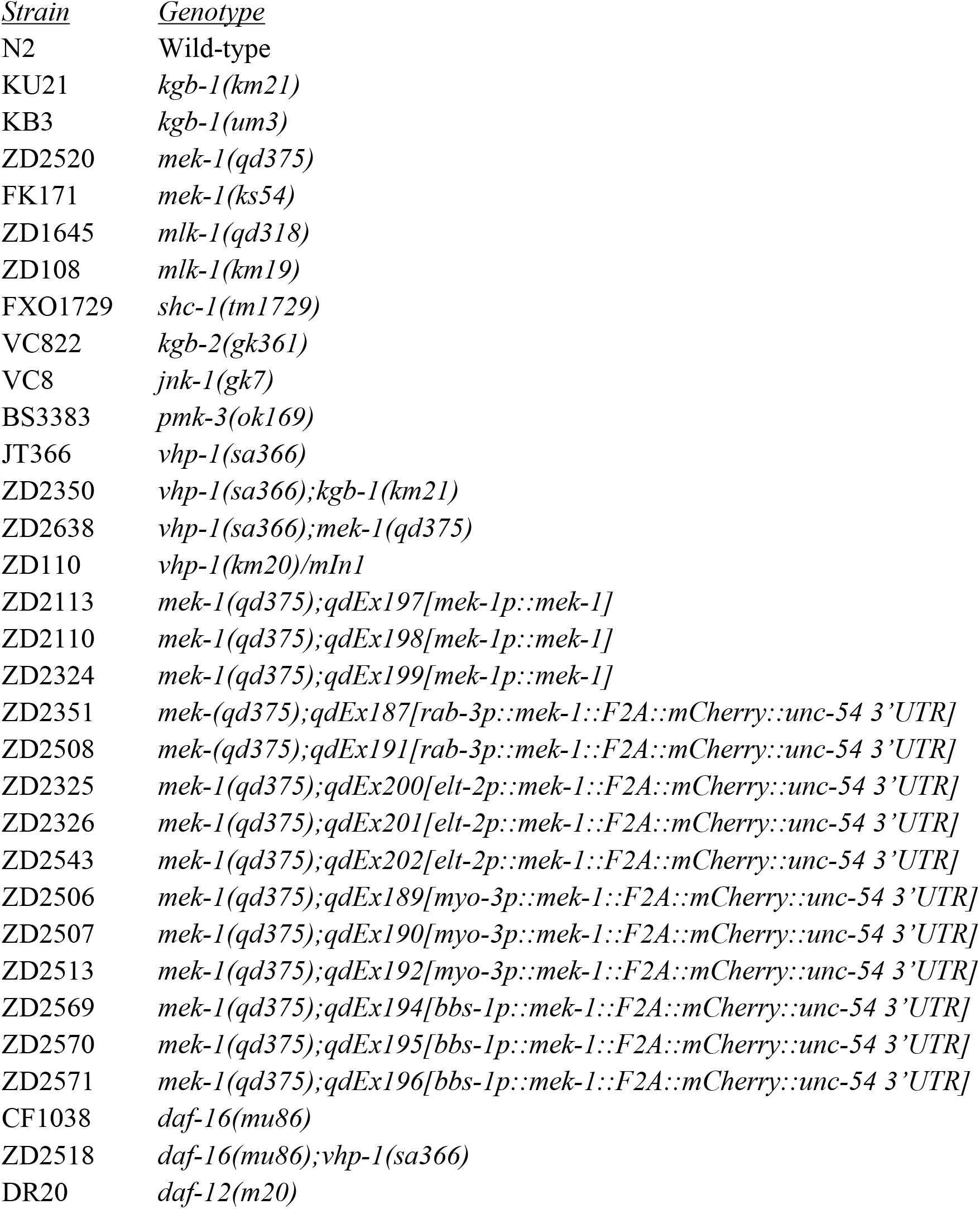
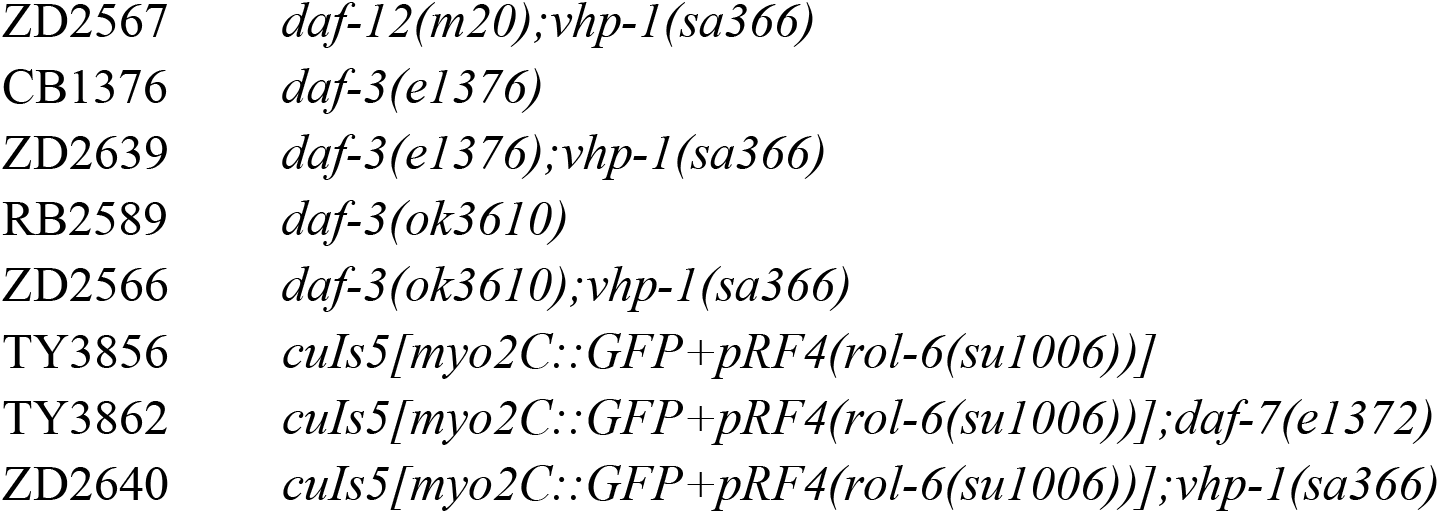

### Dauer formation assays

The dauer assays, unless otherwise mentioned, were performed broadly as described (Neal *et al.* 2013), using the following-indicated pheromone mix, food sources and temperatures. 100 μL of pheromone mix (containing ascarosides ascr#2, ascr#3, ascr#5, and ascr#8 each at a concentration of 20 μM in 10% ethanol) was added to 3.5 cm plates (volume ~ 3 mL) made with Noble agar and without peptone, resulting in an effective plate concentration of 0.67 μM for each ascaroside. The plates were seeded with 20 μl of either concentrated heat-killed (60 mg/mL) or live (70 mg/mL) *E. coli* strain OP50. Three to five gravid animals per individual plate were allowed to lay eggs for three hours at assay temperature (25 °C). The numbers of dauer and non-dauer animals were scored 72 hours (heat-killed food) or 64 hours (live food) post egg-lay midpoint.

### Cloning and generation of transgenic lines

For the genomic rescue construct, the endogenous promoter of *mek-1* (4.2 kb upstream), along with the entire gene and 3’ UTR were amplified from genomic DNA using PCR. This PCR product was cloned into pUC19 backbone using NEBuilder HiFi DNA Assembly (New England Biolabs, Ipswich, MA). The assembled plasmid was microinjected at 25 ng/μl concentration, along with *ofm-1p::gfp* as a co-injection marker at 50 ng/μl for endogenous *mek-1* rescue in the *mek-1* loss-of-function mutant background.

For tissue specific expression constructs, MEK-1 cDNA was amplified from N2 cDNA library using PCR. The cDNA expression was driven by various tissue-specific promoters viz. *elt-2p* (intestine specific expression)*, myo-3p* (body wall muscle specific expression)*, rab-3p* (pan-neuronal expression) and *bbs-1p* (chemosensory neuron specific expression). The promoters along with the cDNA, F2A self-cleaving peptide and mCherry sequence, were all cloned into pPD95.75 plasmid backbone using NEBuilder HiFi DNA Assembly. The mCherry sequence was used to help confirm expression of MEK-1 in the specific tissue and the F2A self-cleaving peptide sequence ensured that they don’t form a fusion protein.

The assembled plasmid for body-wall muscle specific expression was microinjected at 25 ng/μl concentration, along with *ofm-1p::gfp* as a co-injection marker at 50 ng/μl for tissue-specific rescue of *mek-1* in the *mek-1* loss-of-function mutant background. The remaining assembled plasmids were microinjected at 5 ng/μl concentration (due to toxicity issues at higher concentration), along with *ofm-1p::gfp* as a co-injection marker at 100 ng/μl for tissue-specific rescue of *mek-1* in the *mek-1* loss-of-function mutant background.

### Imaging

For measurement of *cuIs5* GFP in the pharynx, animals were mounted with 1 mM sodium azide onto slides with a 2% agarose pad. Slides were viewed using Zeiss AxioImager Z1 fluorescence microscope primarily with 20X objective. Fluorescence signals were recorded with a charge-coupled device camera (AxioCam) using constant exposure time without saturation. Images were captured and processed using AxioVision image processor software.

### Statistical analysis

All statistical analysis was performed using the GraphPad Prism software (Graphpad Prism, RRID:SCR_002798). Statistical tests used are indicated in each figure legend.

## Acknowledgements

We thank the Caenorhabditis Genetics Center, which is funded by NIH Office of Research Infrastructure Programs (P40 OD010440), for providing strains. We thank members of the Kim lab for helpful conversations and feedback on the manuscript and figures. This work is supported by NIGMS grants R01GM084477 and R35GM141794 to D.K. and R35GM131877 to F.S.

## Competing interests

The authors declare that no competing interests exist.

## Notes

### Competing Interest Statement

The authors have declared no competing interest.

## References

1. Golden, J. W., & Riddle, D. L. (1982). A pheromone influences larval development in the nematode *Caenorhabditis elegans*. Science, 218(4572), 578–580.

2. Golden, J. W., & Riddle, D. L. (1984). A *Caenorhabditis elegans* dauer-inducing pheromone and an antagonistic component of the food supply. Journal of chemical ecology, 10(8), 1265–1280.

3. Golden, J. W., & Riddle, D. L. (1984). The *Caenorhabditis elegans* dauer larva: developmental effects of pheromone, food, and temperature. Developmental biology, 102(2), 368–378.

4. Golden, J. W., & Riddle, D. L. (1984). A pheromone-induced developmental switch in *Caenorhabditis elegans*: Temperature-sensitive mutants reveal a wild-type temperature-dependent process. Proceedings of the National Academy of Sciences, 81(3), 819–823.

5. Cassada, R. C., & Russell, R. L. (1975). The dauerlarva, a post-embryonic developmental variant of the nematode *Caenorhabditis elegans*. Developmental biology, 46(2), 326–342.

6. Riddle, D. L., & Albert, P. S. (1997). Genetic and Environmental Regulation of Dauer Larva Development. In C. elegans II, D. L. Riddle, T. Blumenthal, B. J. Meyer, and J. R. Priess, eds. (Plainview, New York: Cold Spring Harbor Laboratory Press), pp. 739–768.

7. Hu, P.J., Dauer (August 08, 2007), WormBook, ed. The C. elegans Research Community, WormBook, doi/10.1895/wormbook.1.144.1, http://www.wormbook.org.

8. Fielenbach, N., & Antebi, A. (2008). *C. elegans* dauer formation and the molecular basis of plasticity. Genes & development, 22(16), 2149–2165.

9. Baugh, L. R., & Hu, P. J. (2020). Starvation Responses Throughout the Caenorhabditis elegans Life Cycle. Genetics, 216(4), 837–878.

10. Butcher, R. A., Fujita, M., Schroeder, F. C., & Clardy, J. (2007). Small-molecule pheromones that control dauer development in *Caenorhabditis elegans*. Nature chemical biology, 3(7), 420–422.

11. Kim, K., Sato, K., Shibuya, M., Zeiger, D. M., Butcher, R. A., Ragains, J. R., Clardy, J., Touhara, K., & Sengupta, P. (2009). Two chemoreceptors mediate developmental effects of dauer pheromone in *C. elegans*. Science, 326(5955), 994–998.

12. McGrath, P. T., Xu, Y., Ailion, M., Garrison, J. L., Butcher, R. A., & Bargmann, C. I. (2011). Parallel evolution of domesticated *Caenorhabditis* species targets pheromone receptor genes. Nature, 477(7364), 321–325.

13. Park, D., O’Doherty, I., Somvanshi, R. K., Bethke, A., Schroeder, F. C., Kumar, U., & Riddle, D. L. (2012). Interaction of structure-specific and promiscuous G-protein–coupled receptors mediates small-molecule signaling in *Caenorhabditis elegans*. Proceedings of the National Academy of Sciences, 109(25), 9917–9922.

14. Larsen, P. L., Albert, P. S., & Riddle, D. L. (1995). Genes that regulate both development and longevity in Caenorhabditis elegans. Genetics, 139(4), 1567–1583.

15. Ren, P., Lim, C. S., Johnsen, R., Albert, P. S., Pilgrim, D., & Riddle, D. L. (1996). Control of *C. elegans* larval development by neuronal expression of a TGF-β homolog. Science, 274(5291), 1389–1391.

16. Schackwitz, W. S., Inoue, T., & Thomas, J. H. (1996). Chemosensory neurons function in parallel to mediate a pheromone response in *C. elegans*. Neuron, 17(4), 719–728.

17. Kimura, K. D., Tissenbaum, H. A., Liu, Y., & Ruvkun, G. (1997). *daf-2*, an insulin receptor-like gene that regulates longevity and diapause in *Caenorhabditis elegans*. Science, 277(5328), 942–946.

18. Kaul, T. K., Rodrigues, P. R., Ogungbe, I. V., Kapahi, P., & Gill, M. S. (2014). Bacterial fatty acids enhance recovery from the dauer larva in Caenorhabditis elegans. PloS one, 9(1), e86979.

19. Khanna, A., Kumar, J., Vargas, M. A., Barrett, L., Katewa, S., Li, P., ... & Kapahi, P. (2016). A genome-wide screen of bacterial mutants that enhance dauer formation in C. elegans. Scientific reports, 6(1), 1–15.

20. Neal, S. J., Takeishi, A., O’Donnell, M. P., Park, J., Hong, M., Butcher, R. A., Kim, K., & Sengupta, P. (2015). Feeding state-dependent regulation of developmental plasticity via CaMKI and neuroendocrine signaling. Elife, 4, e10110.

21. O’Donnell, M. P., Chao, P. H., Kammenga, J. E., & Sengupta, P. (2018). Rictor/TORC2 mediates gut-to-brain signaling in the regulation of phenotypic plasticity in *C. elegans*. PLoS genetics, 14(2), e1007213.

22. Kulalert, W., & Kim, D. H. (2013). The unfolded protein response in a pair of sensory neurons promotes entry of *C. elegans* into dauer diapause. Current Biology, 23(24), 2540–2545.

23. Kulalert, W., Sadeeshkumar, H., Zhang, Y. K., Schroeder, F. C., & Kim, D. H. (2017). Molecular determinants of the regulation of development and metabolism by neuronal eIF2α phosphorylation in *Caenorhabditis elegans*. Genetics, 206(1), 251–263.

24. Malone, E. A., Inoue, T., & Thomas, J. H. (1996). Genetic Analysis of the Roles of *daf-28* and *age-1* in Regulating *Caenorhabditis elegans* Dauer Formation. Genetics, 143(3), 1193–1205.

25. Li, W., Kennedy, S. G., & Ruvkun, G. (2003). *daf-28* encodes a *C. elegans* insulin superfamily member that is regulated by environmental cues and acts in the DAF-2 signaling pathway. Genes & development, 17(7), 844–858.

26. Jeong, P. Y., Jung, M., Yim, Y. H., Kim, H., Park, M., Hong, E., Lee, W., Kim, Y. H., Kim, K., & Paik, Y. K. (2005). Chemical structure and biological activity of the *Caenorhabditis elegans* dauer-inducing pheromone. Nature, 433(7025), 541–545.

27. Butcher, R. A., Ragains, J. R., Kim, E., & Clardy, J. (2008). A potent dauer pheromone component in *Caenorhabditis elegans* that acts synergistically with other components. Proceedings of the National Academy of Sciences, 105(38), 14288–14292.

28. Pungaliya, C., Srinivasan, J., Fox, B. W., Malik, R. U., Ludewig, A. H., Sternberg, P. W., & Schroeder, F. C. (2009). A shortcut to identifying small molecule signals that regulate behavior and development in Caenorhabditis elegans. Proceedings of the National Academy of Sciences, 106(19), 7708–7713.

29. Gallo, M., & Riddle, D. L. (2009). Effects of a *Caenorhabditis elegans* dauer pheromone ascaroside on physiology and signal transduction pathways. Journal of chemical ecology, 35(2), 272–279.

30. Ailion, M., & Thomas, J. H. (2000). Dauer formation induced by high temperatures in *Caenorhabditis elegans*. Genetics, 156(3), 1047–1067.

31. Neal, S. J., Kim, K., & Sengupta, P. (2013). Quantitative assessment of pheromone-induced dauer formation in *Caenorhabditis elegans*. In Pheromone Signaling (pp. 273–283). Humana Press, Totowa, NJ.

32. Sakaguchi, A., Matsumoto, K., & Hisamoto, N. (2004). Roles of MAP kinase cascades in Caenorhabditis elegans. Journal of biochemistry, 136(1), 7–11.

33. Mizuno, T., Hisamoto, N., Terada, T., Kondo, T., Adachi, M., Nishida, E., ... & Matsumoto, K. (2004). The *Caenorhabditis elegans* MAPK phosphatase VHP-1 mediates a novel JNK-like signaling pathway in stress response. The EMBO journal, 23(11), 2226–2234.

34. Mizuno, T., Fujiki, K., Sasakawa, A., Hisamoto, N., & Matsumoto, K. (2008). Role of the Caenorhabditis elegans Shc adaptor protein in the c-Jun N-terminal kinase signaling pathway. Molecular and cellular biology, 28(23), 7041–7049.

35. Nix, P., Hisamoto, N., Matsumoto, K., & Bastiani, M. (2011). Axon regeneration requires coordinate activation of p38 and JNK MAPK pathways. Proceedings of the National Academy of Sciences, 108(26), 10738–10743.

36. Albert, P. S., Brown, S. J., & Riddle, D. L. (1981). Sensory control of dauer larva formation in Caenorhabditis elegans. Journal of Comparative Neurology, 198(3), 435–451.

37. Bargmann, C. I., & Horvitz, H. R. (1991). Control of larval development by chemosensory neurons in *Caenorhabditis elegans*. Science, 251(4998), 1243–1246.

38. Bargmann, C.I. Chemosensation in *C. elegans* (October 25, 2006), WormBook, ed. The C. elegans Research Community, WormBook, doi/10.1895/wormbook.1.123.1, http://www.wormbook.org.

39. Ludewig AH. and Schroeder FC. Ascaroside signaling in *C. elegans* (January 18, 2013), WormBook, ed. The C. elegans Research Community, WormBook, doi/10.1895/wormbook.1.155.1, http://www.wormbook.org.

40. Choy, R. K., & Thomas, J. H. (1999). Fluoxetine-resistant mutants in C. elegans define a novel family of transmembrane proteins. Molecular cell, 4(2), 143–152.

41. Vowels, J. J., & Thomas, J. H. (1992). Genetic analysis of chemosensory control of dauer formation in *Caenorhabditis elegans*. Genetics, 130(1), 105–123.

42. Thomas, J. H., Birnby, D. A., & Vowels, J. J. (1993). Evidence for parallel processing of sensory information controlling dauer formation in *Caenorhabditis elegans*. Genetics, 134(4), 1105–1117.

43. Thatcher, J. D., Haun, C., & Okkema, P. G. (1999). The DAF-3 Smad binds DNA and represses gene expression in the *Caenorhabditis elegans* pharynx. Development, 126(1), 97–107.

44. Reiner, D. J., Ailion, M., Thomas, J. H., & Meyer, B. J. (2008). *C. elegans* anaplastic lymphoma kinase ortholog SCD-2 controls dauer formation by modulating TGF-β signaling. Current Biology, 18(15), 1101–1109.

45. Koga, M., Zwaal, R., Guan, K. L., Avery, L., & Ohshima, Y. (2000). A Caenorhabditis elegans MAP kinase kinase, MEK-1, is involved in stress responses. The EMBO journal, 19(19), 5148–5156.

46. Uno, M., Honjoh, S., Matsuda, M., Hoshikawa, H., Kishimoto, S., Yamamoto, T., ... & Nishida, E. (2013). A fasting-responsive signaling pathway that extends life span in *C. elegans*. Cell reports, 3(1), 79–91.

47. Kim, D. H., Liberati, N. T., Mizuno, T., Inoue, H., Hisamoto, N., Matsumoto, K., & Ausubel, F. M. (2004). Integration of *Caenorhabditis elegans* MAPK pathways mediating immunity and stress resistance by MEK-1 MAPK kinase and VHP-1 MAPK phosphatase. Proceedings of the National Academy of Sciences, 101(30), 10990–10994.

48. Huffman, D. L., Abrami, L., Sasik, R., Corbeil, J., van der Goot, F. G., & Aroian, R. V. (2004). Mitogen-activated protein kinase pathways defend against bacterial pore-forming toxins. Proceedings of the National Academy of Sciences, 101(30), 10995–11000.

49. Twumasi-Boateng, K., Wang, T. W., Tsai, L., Lee, K. H., Salehpour, A., Bhat, S., ... & Shapira, M. (2012). An age-dependent reversal in the protective capacities of JNK signaling shortens *Caenorhabditis elegans* lifespan. Aging cell, 11(4), 659–667.

50. Liu, L., Ruediger, C., & Shapira, M. (2018). Integration of stress signaling in *Caenorhabditis elegans* through cell-nonautonomous contributions of the JNK homolog KGB-1. Genetics, 210(4), 1317–1328.

51. Aghayeva, U., Bhattacharya, A., Sural, S., Jaeger, E., Churgin, M., Fang-Yen, C., & Hobert, O. (2021). DAF-16/FoxO and DAF-12/VDR control cellular plasticity both cell-autonomously and via interorgan signaling. PLoS biology, 19(4), e3001204.

52. Neal, S. J., Park, J., DiTirro, D., Yoon, J., Shibuya, M., Choi, W., ... & Sengupta, P. (2016). A forward genetic screen for molecules involved in pheromone-induced dauer formation in *Caenorhabditis elegans*. G3: Genes, Genomes, Genetics, 6(5), 1475–1487.

